# Comparison of AI protein structure ensemble prediction tools

**DOI:** 10.64898/2026.05.29.728804

**Authors:** Lisa Otten, Jeremy M. G. Leung, Lillian T. Chong, Daniel M. Zuckerman

## Abstract

Multiple AI prediction tools for protein structural ensembles have recently been released, building on the much heralded advances from AlphaFold, large language models, and other machine-learning approaches. Here we report on a comparison of several tools (BioEmu, AFSample2, ESMFlow) using a small test set of proteins, including three which exhibit well-studied structural transitions. Overall, while the AI platforms generate structurally diverse ensembles with overlapping regions, each tool produces clearly distinct conformational distributions. Thus, it is impossible that all the tools generate ensembles of high biophysical quality, analogous to a Boltzmann distribution. Experimental structures are often, but not always, covered by the ensembles in dimensionally reduced spaces. In cases where point mutations are known experimentally to cause large structural shifts, the AI tools exhibit either small or negligible shifts. Although our current analysis cannot evaluate the absolute quality of an ensemble, and hence cannot identify a best-performing AI tool, the results suggest users pursuing downstream applications such as protein engineering or drug design should interpret these ensembles with caution.

## Introduction

The field of molecular simulation is predicated on the now widely accepted paradigm that an individual protein structure generally is insufficient to provide a complete characterization of a protein’s function.^1-4^ The ideal description would be a time-ordered sequence of structures of the protein performing its function—in fact, the true ideal would be many such trajectories to account for stochastic variability in the dynamics.^5^ Because the true temporal dynamics typically are out of reach both experimentally and computationally, considerable attention has been focused on the (non-temporal) ensemble description of structural fluctuations.^6-9^ In the case of equilibrium, the ensemble of structures should be distributed according to the Boltzmann factor, although nonequilibrium steady-state ensembles are also highly pertinent for biology.^10,11^

Building on the success of machine learning and “artificial intelligence” (AI) methods for the prediction of individual protein structures,^12-16^ there has been a recent spurt of development of tools for ensemble generation. Our study focuses on three tools: (i) AFSample2^17^ which manipulates the multiple-sequence alignment step in AlphaFold2 to reduce co-evolution information and thus generate a diversity of structure predictions, (ii) BioEmu^18^ which also uses AlphaFold methods, augmented by a diffusion model and training on extensive molecular dynamics simulation data, and (iii) ESMFlow^19^ which employs large-language model transformer methodology. Our study examines two ESMFlow models, both trained on experimental Protein Data Bank (PDB) structures but one of them further fine-tuned on simulated molecular dynamics (MD) trajectories.

How should AI ensemble generation tools be assessed? On the one hand, a definite goal is to improve the “coverage” of conformational states to ensure better prediction of “alternative states”.^17^ On the other hand, the biophysical goal of producing “equilibrium ensembles” properly distributed according to the Boltzmann factor^18^ is a higher standard. In this study, we examine both coverage and distribution, although as detailed below, we cannot directly evaluate the distribution of a single ensemble.

Our approach is to compare the coverage and distributions of the different tools, which is done by comparing outputs of the tools to one another and by comparison to known experimental structures. We attempt to provide a detailed analysis of a small set of four proteins. Three of these exhibit significant, experimentally documented conformational motions: adenylate kinase (ADK), calmodulin (CaM), and dihydrofolate reductase (DHFR). The fourth is a CATH domain which also has several experimental structures.

Overall, the data below suggest the AI tools likely have not reached an ideal level of performance. There is minimal consensus among predictions generated by the AI tools, both in terms of coverage and distribution. There is particular ambiguity surrounding binding: the tools are trained on both liganded and ligand-free structures, but the two forms don’t necessarily co-exist significantly in all binding states and the AI tools examined do not allow specification of binding state. Further, the predictions are relatively insensitive to point mutations known to cause significant conformational changes. The current analysis, because of a lack of ground truth, cannot choose a “winner” or endorse any particular AI tool. We do believe the path forward is to rely on physics-based sampling and reweighting methods^20^ to refine AI ensembles which currently are unverifiable and ostensibly less than fully reliable.

## Results

We discuss our findings system by system, providing background on each protein to help contextualize the results. Note that all the tools have been trained on the full Protein Data Bank (PDB), i.e., including the very experimental structures being compared against below. ESMFlow-MD is an extension of the ESM-PDB model, further fine-tuned on explicitly solvated, all-atom molecular dynamics simulations from the ATLAS database.^21^ On the whole, our expectation for coverage of experimental structures should be high because of the training data. Note that because BioEmu does not generate side-chain coordinates, those have been supplied by the H-Packer tool^22^ as recommended by BioEmu—see Methods. It is also worth noting that experimental structures reflect their own limitations: crystallization conditions in the case of X-ray structures; both the modeling protocol and solution conditions in the case of NMR. It is fair to say there is no physiological ground truth available, though comparison with experimental structures remains informative.

### Adenylate Kinase (ADK) - Background

ADK is known to exhibit significant structural variability, notably open and closed forms differing by 7 Å RMSD in the bacterial form.^23^ For human ADK examined here (Fig. 1), structures of ligand-bound states are all closed, as are all unliganded (apo) wild-type structures, but two unliganded point mutants (R97W) and (R128W) experimentally exhibit open states. However, the ADK sequences and structures most commonly used for MD simulations are from bacteria, where the unliganded form is open.^23,24^ ADK may exhibit some population of each state in both apo and liganded states, but the populations of the states are unknown in the different conditions.

**Fig. 1.**
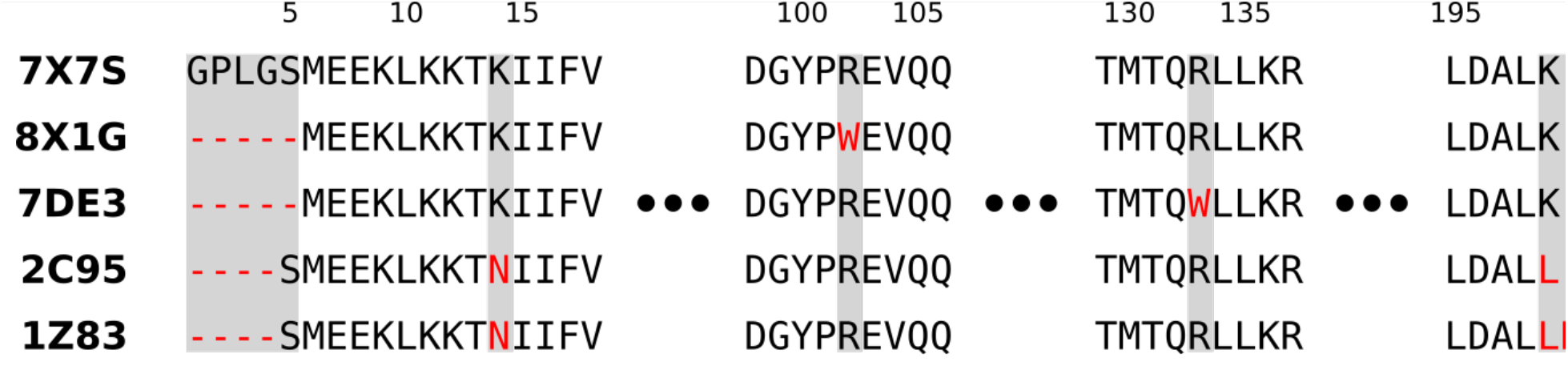
Sequence alignment of five adenylate kinase PDB structures. Residue sequence of the five PDB structures shown in Fig. 2. The first sequence (7 × 7S) is used in the AI tools. Differences in sequences are marked (red, gray). Includes unpublished information from https://doi.org/10.2210/pdb7DE3/pdb and https://doi.org/10.2210/pdb8×1G/pdb.

The ligand binding state of ADK cannot be specified in the AI tools examined which creates a fundamental ambiguity: does the ensemble represent the apo state, a liganded state, or some unphysical mixture? ADK catalyzes the reaction ATP + ADP ⇔ 2 ADP and thus in principle can bind either ‘reactants’ or ‘products’, as well as other molecules, notably chimeras that do not undergo catalysis.^25^ The AI tools are trained on both apo and liganded structures without distinction, and thus it is unclear whether the ensembles they generate should correspond to any precise biophysical state.

### Adenylate Kinase - Results

Fig. 2 shows an analysis of AI ensembles for ADK considering both backbone and sidechain behavior. We consider (i) the issue of ‘coverage’, i.e., whether the conformation space of the ensembles includes known experimental structures for the same (human) isoform of ADK, (ii) the consistency among the AI ensembles, and (iii) the sensitivity of AI tools to point mutations. Current data analysis is largely based on principal components analysis (PCA), which enables visualization of the ensembles across the most variable PCs. PCA is applied to both the protein backbone atomic coordinates and separately to side-chain chi angles as described in Methods, in both cases based on a joint ensemble aggregated from all the AI tools.

**Fig. 2.**
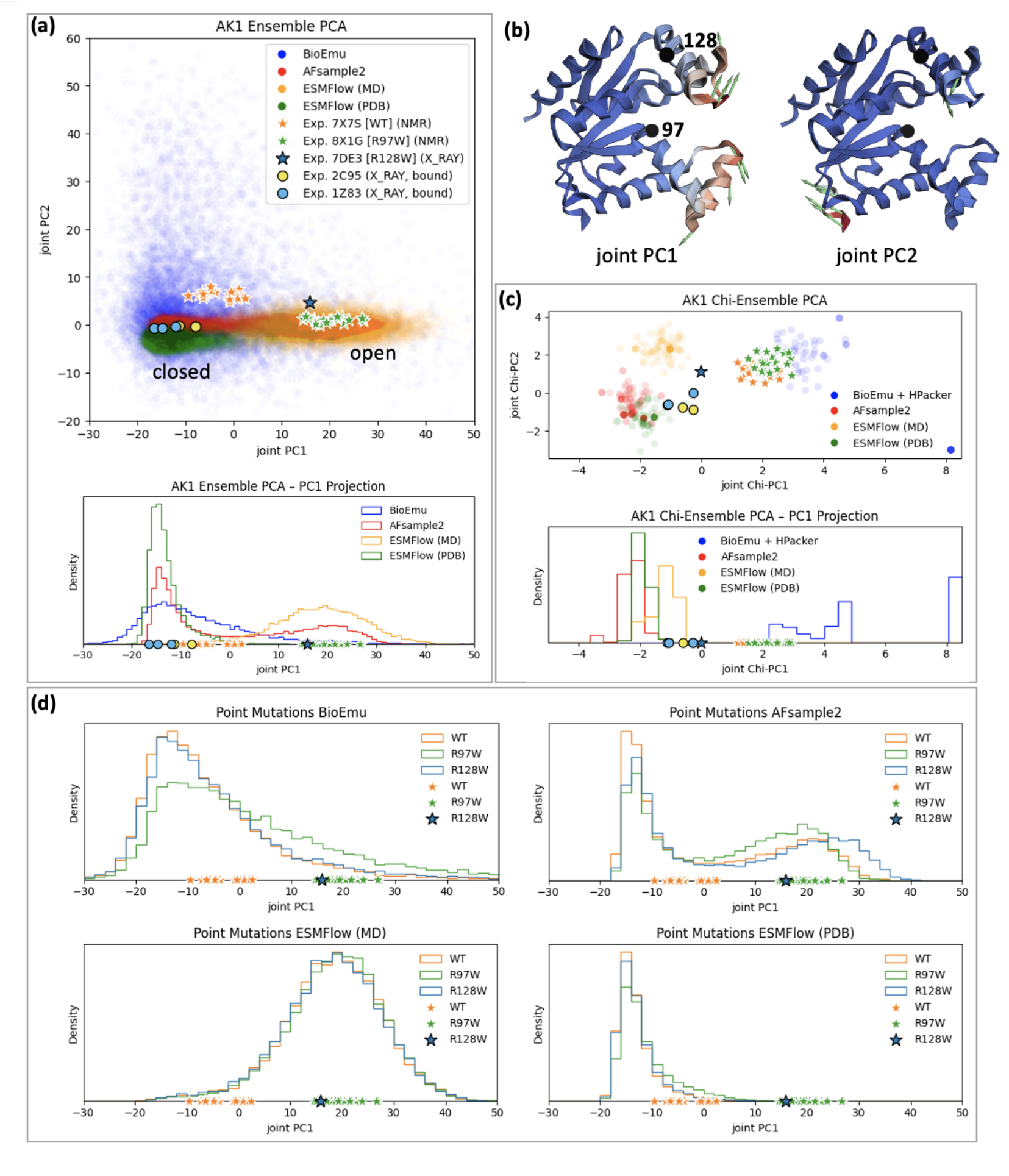
Adenylate kinase conformational ensembles from AI tools. (a) Protein backbone structures projected onto principal components (PCs) constructed from the aggregated ensembles. Also shown are experimental structures, with ligand-bound forms indicated by circles and unliganded (apo) forms indicated by stars. NMR structures are outlined in white, whereas crystal structures are outlined in black. (b) Structures illustrating the first two backbone PCs, with redder colors depicting greater participation and arrows depicting the dominant direction of variability. (c) Projection of subsampled ensembles (∼10,000 structures subsampled to 42-62 structures, see Methods) onto PCs defined based on the joint distribution of side-chain chi angle sines or cosines. Note that BioEmu structures lack side chains, which were modeled here using H-Packer. (d) Predictions of AI tools for R97W and R128W point mutations, as compared to wild-type (WT). Distributions shown in the space of the joint backbone PC1 as in (a).

#### Backbone coverage and consistency

We first consider coverage of experimental structures in the space of backbone PCs as shown in Fig. 2(a). Because the experimental structures represent highly distinct conditions, as noted above, it is unclear whether it is desirable to cover all these structures or just a subset appropriate to one condition (e.g., apo). BioEmu’s structures are overwhelmingly closed and exhibit the broadest spread in PC1-PC2 space, which is largely because other tools do not exhibit substantial variation in the PC2 direction. AFSample2 exhibits the only bimodal distribution, covering both closed and open conformations without reaching some of the NMR structures which are slightly displaced in PC2. The two versions of ESMFlow exhibit opposite tendencies toward open vs. closed structures and also do not cover the same NMR structures missed by AFSample2. Lacking ground truth, it is impossible to say one tool is better than the other, and as noted above, all the tools were explicitly or implicitly trained on PDB data.

#### Sidechain coverage and consistency

Fig. 2(c) illustrates the distributions of sidechain chi angles generated by AI tools, along with experimental structures. Note that because BioEmu does not generate sidechains, those have been supplied by the H-Packer tool.^22^ Among the AI ensembles, AFSample2 and ESMFlow-PDB show overlap in the PC1-PC2 space with NMR ensembles, although not with X-ray structures. ESMFlow shows no overlap with the experimental structures and BioEmu+H-Packer is yet more distant. Note that side-chain analysis for all proteins was performed with a downsampled set of structures – see Methods.

#### Sensitivity to point mutations

Fig. 2(d) depicts the extent to which point mutations shift the ensembles of adenylate kinase predicted by each AI tool. While the ground truth is unknown, available experimental structures suggest both mutants should produce significant rightward shifts along PC1. In practice, all of the AI tools exhibit relatively modest shifts. For the R97W mutation, BioEmu, AFSample2, and ESMFlow-PDB each show a visible rightward shift. For R128W, only AFSample2 produces a noticeable shift. ESMFlow-MD exhibits negligible sensitivity to either mutation.

### CATH Domain 2wg5F02 - Background

CATH domains derive from a hierarchical organization of PDB structures according to Class(C), Architecture(A), Topology(T) and Homologous superfamily (H).^26^ CATH domains provide representative structures common to numerous (often, multi-domain) proteins, thus providing an organizing basis for structural biology. CATH domains were a key focus of the recent BioEmu paper,^18^ and here we select one of their best-performing examples from that study.

### CATH Domain 2wg5F02 - Results

#### Backbone Coverage and Consistency

**Fig. 3(a)** shows AI ensembles based on the protein backbone which exhibit excellent overlap with the experimental structures, arguably for all the AI ensembles although some structures are in the tail of the AFSample2 distribution. The AI ensembles all overlap with one another in the central values of the PCs but have substantial differences in width as well as shifts in their mean values.

**Fig. 3.**
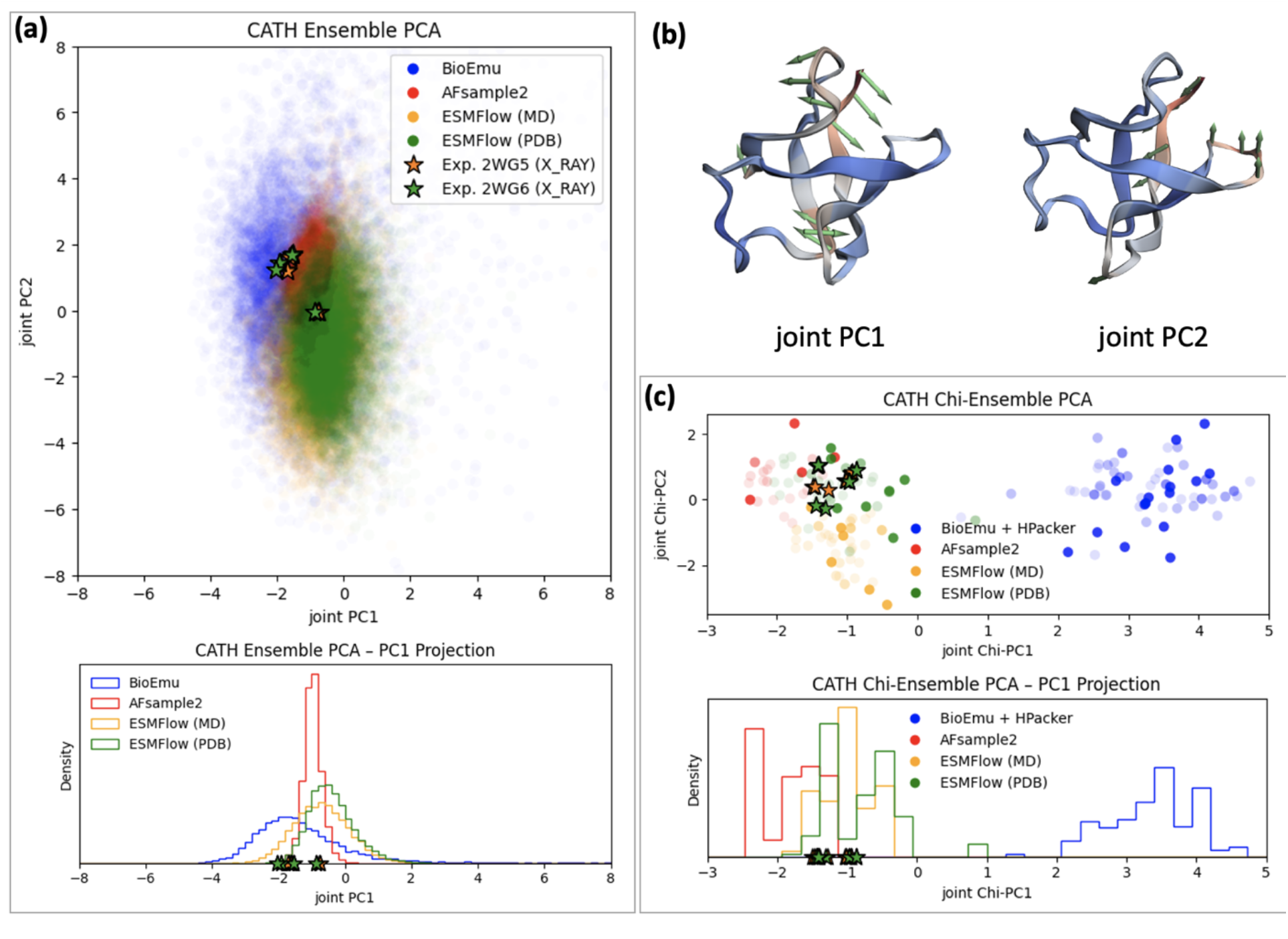
CATH domain 2wg5F02 conforma5onal ensembles from AI tools. (a) Protein backbone structures projected onto principal components (PCs) constructed from the aggregated ensembles. Also shown are experimental structures, with ligand-bound forms indicated by circles and unliganded (apo) forms indicated by stars. NMR structures are outlined in white, whereas crystal structures are outlined in black. (b) Structures illustrating the first two backbone PCs, with redder colors depicting greater participation and arrows depicting the dominant direction of variability. (c) Projection of ensembles (∼10,000 structures subsampled to 21-62 structures, see Methods) onto PCs defined based on the joint distribution of side-chain chi angle cosines. Note that BioEmu structures lack side chains, which were modeled here using H-Packer.

#### Sidechain coverage and consistency

In the space of sidechain PCs (**Fig. 3(c)**), all the ensembles except for BioEmu cover the at least some of the values of the experimental structures, and we note again that BioEmu sidechains are generated by H-Packer. The sidechain distributions are also shifted from one another: AFSample2 is shifted from both ESMFlow models, with BioEmu being disjoint based on the subsampled ensembles shown.

No point mutations are analyzed for the CATH domain.

### Calmodulin (CaM) C-terminal domain – Background

Calmodulin is a two-domain calcium sensor, undergoing conformational changes that regulate its interactions with a wide range of protein targets.^27-29^ For simplicity and more facile visualization of the results, we examine solely the C-terminal domain, building on prior simulation work examining one domain at a time.^30,31^ Because most experimental structures were determined with calcium bound, we anticipate that AI tools will implicitly model a calcium-bound condition.

### Calmodulin C-terminal domain - Results

#### Backbone Coverage and Consistency

From the one and two-dimensional distributions depicted in **Fig. 4(a)**, we see that all of the experimental structures are covered by at least one of the AI tools in the space of the first two PCs (**Fig. 4(b)**). Note that, among experimental structures, only the NMR structures 1CFC and 1CFD are both calcium free and unliganded, leading to the most open structure; all the other structures are bound to a peptide ligand. AFSample2 exhibits the narrowest distribution and hence fails to cover the most experimental structures, followed by ESMFlow-PDB. On the whole, the distributions along PC1, which quantifies opening-closing among the two helix pairs, are highly distinct among the AI tools. Once again, BioEmu exhibits by far the broadest distribution, and as with adenylate kinase (but not the CATH domain) exhibits by far the widest variation in PC2.

**Fig. 4.**
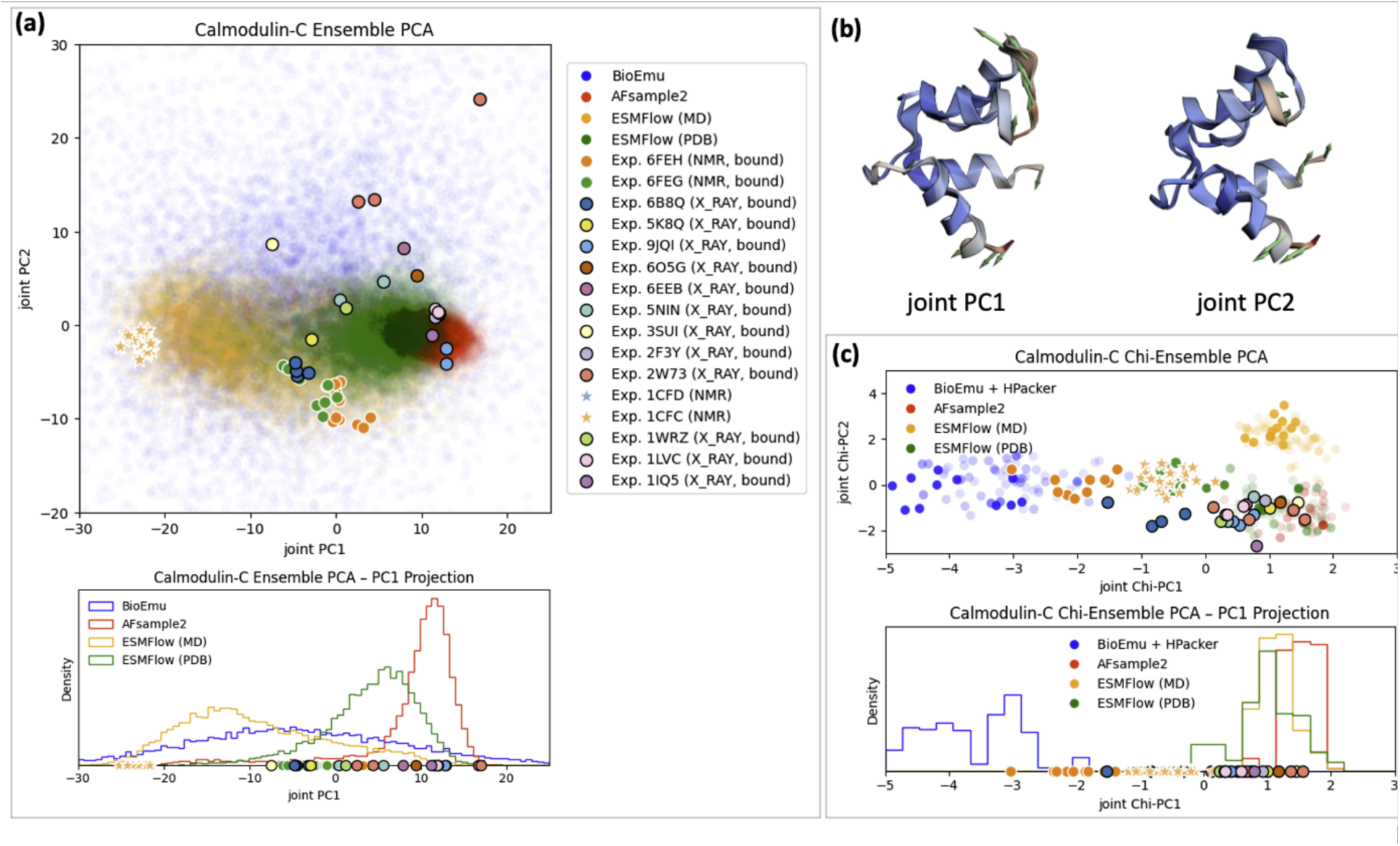
Calmodulin C-terminal domain conforma5onal ensembles from AI tools. (a) Protein backbone structures projected onto principal components (PCs) constructed from the aggregated ensembles. Also shown are experimental structures, with ligand-bound forms indicated by circles and unliganded (apo) forms indicated by stars. NMR structures are outlined in white, whereas crystal structures are outlined in black. (b) Structures illustrating the first two backbone PCs, with redder colors depicting greater participation and arrows depicting the dominant direction of variability. (c) Projection of ensembles (subsampled to 38-74 structures) onto PCs defined based on the joint distribution of side-chain chi angle cosines. Note that BioEmu structures lack side chains, which were modeled here using H-Packer.

#### Sidechain coverage and consistency

The sidechain comparison (**Fig. 4(c)**) matches the behaviors seen in other proteins. The BioEmu/H-Packer sidechains are largely outliers compared to the experimental and AI structures, although there is some overlap in the right tail of the first PC. ESMFlow-MD overlaps with experimental structures in PC1 but not PC2, whereas AFSample2 and ESMFlow-PDB exhibit the best coverage of experimental structures.

### Dihydrofolate Reductase (DHFR) – Background

DHFR is an essential metabolic enzyme that catalyzes the NADPH-dependent reduction of dihydrofolate to tetrahydrofolate, a key reaction required for thymidylate, purine, and amino acid biosynthesis and therefore critical for cellular proliferation.^32^ Beyond its metabolic role, DHFR, and particularly that from Escherichia coli, has become a paradigmatic system for investigating the interplay between protein conformational dynamics and enzymatic catalysis. Extensive structural, kinetic, NMR, and computational studies have shown that DHFR samples a network of interconverting conformational states during its catalytic cycle, including closed, occluded, and open substates associated with substrate binding, hydride transfer, and product release.^33-35^

### DHFR – Results

#### Backbone Coverage and Consistency

The one and two-dimensional distributions depicted in **Fig. 5(a)** reveal that the AI-generated ensembles sample only a subset of the experimentally observed conformational space along the first principal component (**Fig. 5(b)**). Note that all experimental structures shown correspond to liganded structures with diverse ligands. The structures generated by the AI tools vary mostly along PC1 with AFSample2 exhibiting the narrowest distribution, although it still overlaps with several experimental structures. In contrast, ESMFlow-MD samples one of the broadest PC1 distributions, comparable to BioEmu, yet shows minimal overlap with the experimental structures. BioEmu provides the broadest experimentally relevant coverage, capturing the largest fraction of the observed structural states. This set of ensembles shows that broad conformational diversity alone does not necessarily translate into experimentally relevant sampling.

**Fig. 5.**
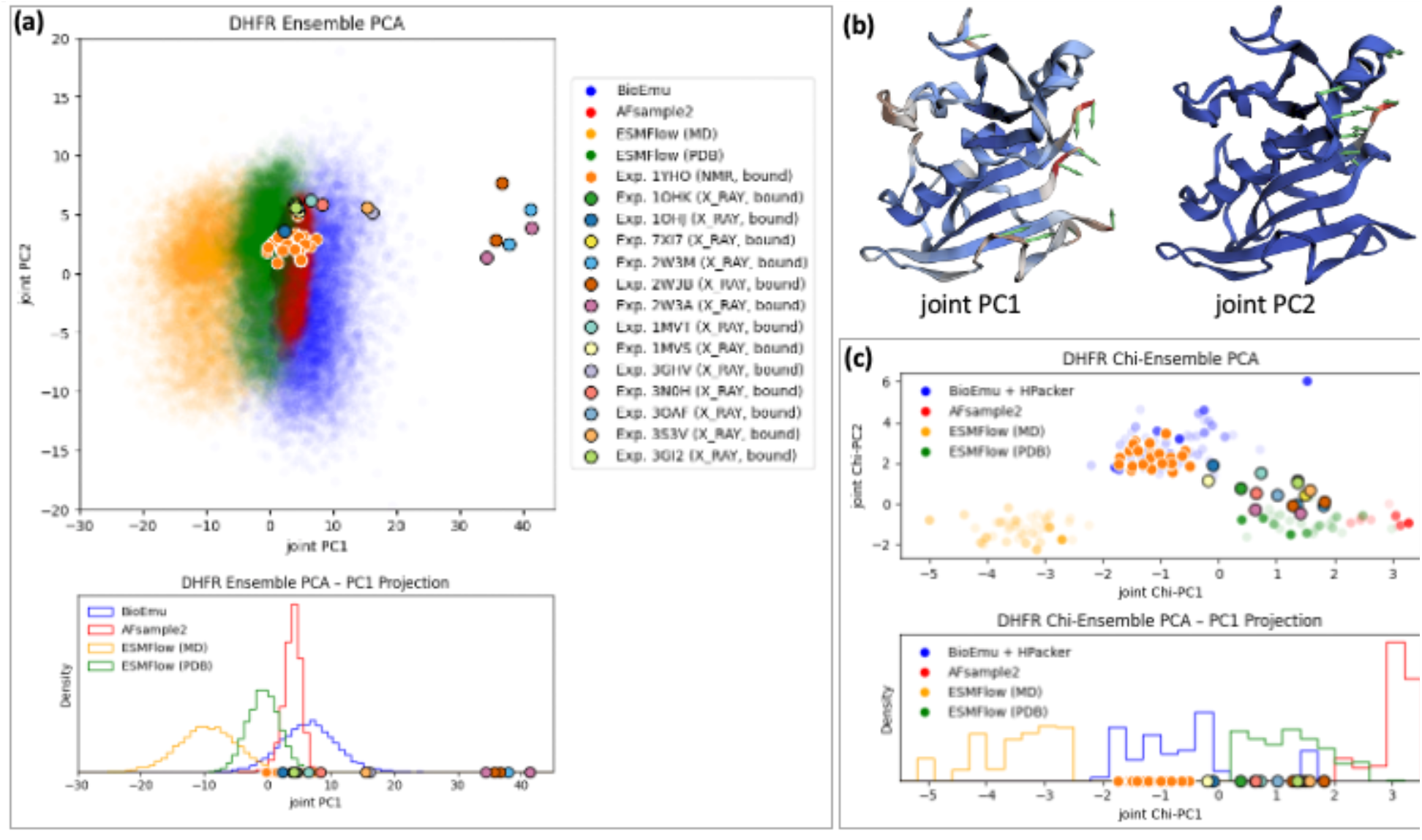
Dihydrofolate reductase conformational ensembles from AI tools. (a) Protein backbone structures projected onto principal components (PCs) constructed from the aggregated ensembles. Also shown are experimental structures, with ligand-bound forms indicated by circles and unliganded (apo) forms indicated by stars. NMR structures are outlined in white, whereas crystal structures are outlined in black. (b) Structures illustrating the first two backbone PCs, with redder colors depicting greater participation and arrows depicting. (c) Projection of ensembles (subsampled to 12-55 structures) onto PCs defined based on the joint distribution of side-chain chi angle cosines. Note that BioEmu structures lack side chains, which were modeled here using H-Packer.

#### Sidechain coverage and consistency

As for previous proteins, the sidechain comparison (**Fig. 5(c)**) shows that the BioEmu/H-Packer sidechains match best with the experimental structures generated by NMR. ESMFlow-MD sidechains are largely outliers, especially along Chi-PC1, while ESMFlow-PDB exhibits the best coverage of experimental structures in the first two PC’s.

## Concluding Discussion

As the roles and prominence of AI tools in research continue to grow, these tools merit an accordingly higher level of scrutiny. Because of their vastly improved speed and ease-of-use compared to conventional MD-based computations, the appeal of such tools is undeniable. We believe the AI tools are here to stay and are worth leveraging. We also believe that the community will benefit from continual evaluation of the tools.

When it comes to protein structural ensembles, absolute ground truth is lacking, but we have pursued meaningful assessment through two approaches: (i) comparing the outputs of different AI tools to one another, and (ii) comparing AI outputs to experimental structures. We do not consider experimental structures as ground truth for equilibrium ensembles because the structures come with important caveats, such as unphysiological experimental conditions and the assumptions built into computational modeling of raw data. We note that some experimental methods, such as room-temperature crystallography,^36^ DEER EPR,^37^ and ^19^F NMR,^38^ can provide direct structural information and state population data, and benchmarking against such data would be valuable future direction. In ongoing work described elsewhere,^20^ we are also comparing AI ensembles to physics-based sampling, which is subject to the usual limitations of sampling convergence and force field accuracy.

Several results stand out from our analysis of ensembles across multiple proteins using four AI tools. Most importantly, the tools produce inconsistent results, both for backbone and side-chain degrees of freedom. Thus, we can be certain that all tools are not giving the (unique) equilibrium ensemble, but we cannot rule out that one tool is truly on target. We also found that none of the tools exhibits the degree of sensitivity to point mutations expected from experiment, though the experimental caveats noted above apply here as well. In many cases, the tools generated ensembles encompassing structures similar to known experimental structures, though this is not surprising given that the AI models were trained on structural data. A notable exception is DHFR, where all of the AI tools failed to generate structures resembling an important subset of experimental structures.

Because the field of ensemble prediction is evolving rapidly, we anticipate these results will need to be revisited with regularity. Nevertheless, we hope our initial findings will be of immediate value and the proposed analyses should be useful as the field develops.

## Methods

### Backbone analysis

All AI tools were configured to generate 10,000 structures for backbone analysis using default/suggested settings.

### Preparation of a “downsampled” ensemble from the full AI-generated ensemble

Side-chain analysis was performed using downsampled ensembles to save computational cost when using H-Packer.

Downsampling was performed using a sequence of steps. To identify a reference structure for alignment, the AI-generated ensembles were first clustered using DBSCAN on pairwise Cα distance matrices computed using the CPPTRAJ program in the Amber software package. Optimal DBSCAN parameters were selected using the kdist “elbow” method for k=2 to k=10. In cases where more than one cluster was identified, the center of the largest cluster was used as the alignment reference; no secondary clusters contained more than 1% of the dataset. Principal component analysis (PCA) was then performed on the aligned Cα coordinates. The AFSample2 and ESMFlow ensembles were clustered jointly to define a shared reference structure and PC space, whiles the BioEmu ensembles were analyzed separately.

Each AI ensemble was projected onto the first two dimensions of its corresponding PC space, and PC1 and PC2 were each divided into 10 equally spaced bins spanning their respective minimum and maximum values. The downsampled ensemble was constructed by randomly selecting one structure from each occupied bin, with each selected structure assigned a weight proportional to the number of structures in each bin. Missing atoms and sidechains were reconstructed as needed.

The resulting 20-80 structures were then energy-minimized and equilibrated using the Amber ff14SB-onlysc force field and a generalized Born implicit solvent model (GBNeck2)^39,40^ at 298K and a water-like collision frequency γ of 80 ps^-1^. Each structure was subjected to 1000 steps of unrestrained minimization (500 steepest descent, 500 conjugate descent), 50 ps of backbone-restrained NVT equilibration (1 kcal mol^-1^A^-2^), and 1 ps of unrestrained NVT equilibration.

### Sidechain reconstruction for the BioEmu-generated ensemble

Because BioEmu generates only backbone coordinates, sidechains were reconstructed using the H-Packer tool,^22^ as recommended by BioEmu, with default settings. These settings included a two-stage equilibration of the reconstructed structures using the OpenMM simulation software with the Amber ff99SB force field, TIP3P explicit solvent, and harmonic positional restraints applied to backbone heavy atoms throughout: 100 ps at constant temperature (300K), followed by 400 ps at constant temperature (300K) and pressure (1 atm).

## Acknowledgements

LTC and DMZ gratefully acknowledge support from the NIH via Grant GM115805. The research reported in this publication used computational infrastructure supported by the OHSU Office of Research Infrastructure Programs, Office of the Director, of the NIH under Award No. S10OD034224.

